# Calories delivered by nonhuman primate foraging enrichment: Data and methods to inform decisions concerning the dietary consequences of enrichment

**DOI:** 10.1101/2020.05.20.106781

**Authors:** Peter J. Pierre, Nicolette A. Torres, Michele Rosga, Jennifer Sullivan

## Abstract

Foraging opportunities are a key component of enrichment in captive nonhuman primates providing manipulative opportunities in which animals can engage in species-typical behaviors. Recent studies suggest captive NHP populations have increased body weight over time leading to negative health outcomes. Increasing food foraging opportunities provides added calories that can be estimated but often are not directly measured. We assessed 10 commonly used foraging puzzles for the amount of foraging material they held; evaluated the range of caloric content delivered; and determine the labor and usage cost of each. Five preparation areas were sampled. The amount of foraging material contained in each prepared puzzle was calculated by subtracting the average empty puzzle weight from the loaded puzzle weight. A detailed description of food content was used to illustrate caloric content. Larger prepared puzzles held more food (M=104.9 g) as compared to small volumes puzzles (M=59.8 g). Analysis of the weight of food applied and caloric content suggest larger puzzles do have the potential to provide increased caloric load; however, the added food from large foraging puzzles constitutes only 5%-10% of the daily k/cal dietary requirement. The study illustrates the importance of considering puzzle characteristics when selecting foraging toys and discusses the considerations of cost of puzzle deployment, maintenance and interactive benefit. The broad significance of our assessment suggests that calories from foraging opportunities are not excessive and can be managed by adopting standard serving sizes and scheduling intermitted presentation of non-nutritive enrichment options.

## Introduction

Environmental enrichment (EE) is a central part of best practices in care for nonhuman primates in the range of captive settings that include dedicated research facilities. The importance of EE to those charged with care of nonhuman primates, as well as to broader public and regulatory agencies, is reflected in its specific provision within federal law. Over 30 years ago, the US Animal Welfare Act (AWA) was amended to require formal environmental enrichment plans for all facilities that house captive primates and that are registered with the United States Department of Agriculture for research or are licensed by the USDA as exhibitors, dealers, or breeders (Public Law 99-158. 1985). For research facilities, the AWA requirement was carried forward and expanded upon in the National Research Council’s publication, “The Guide for the Care and Use of Laboratory Animals” (National Research Council. 1996; National Research Council. 2011).

Efforts to enrich the environments in which nonhuman primates live in captivity have increased over time. Those efforts are generally agreed to be worthwhile and to result in positive outcomes for animals’ health and well-being. Nonetheless, implementation and maintenance of environmental enrichment programs pose inherent challenges. Among the challenges to selection and implementation of enrichment is the effect of enrichment on animals’ health and on scientific objectives. The empirical literature highlights both the potential positive and negative effects of enrichment, and the challenges to evaluating what constitutes increased well-being (Novak & Suomi,1988; Wolfle, 1999). For example, Bayne (2005) points out that enrichment cannot be considered only in terms of benefit, but that physical harm and deleterious changes in animals’ physiological responses can also occur as a result of implementing enrichment strategies. Similarly, Nelson and Mandrell (2005), outline six premises for consideration of enrichment strategies, highlighting both positive and negative indicators of their relative effectiveness. In a specific example, a recent study reviewed the potential for nutritive elements of food enrichment to have negative effects that can interfere with outcome measures in toxicology studies, and, in turn, jeopardize the scientific objectives for which the animals are maintained in the research setting (Cooper, 2015).

The use of food in environmental enrichment is ubiquitous and there is general consensus that food items should be comprised of “healthy dietary choices”. It is, however, not well studied, tends to vary widely in practice, and often raises concerns in terms of animal health and interference with scientific objectives. Furthermore, numerous stakeholders with a diverse set of specialty expertise, job functions, and interests, that include: Scientists, funding agencies, facility managers, veterinarians, and care staff have input to the “Enrichment” process (Bennett, et al., 2010; 2014; 2016). The concerns of these stakeholders range from animal welfare to scientific objectives to costs, labor, facility maintenance and procedural compliance.

Our broad goal is to promote evidence-based decision-making. The current study illustrates a method that can be adopted to inform decisions, facilitate record keeping and allow for direct comparisons between enrichment strategies and puzzles, across time, and across facilities. We focus on a method developed to provide quantitative assessment of one type of enrichment, foraging puzzles, to provide information that could play a role in decisions about nonhuman primate environmental enrichment.

### Foraging opportunities as nonhuman primate environmental enrichment

Free ranging macaques spend approximately 40% of the day foraging (Goldstein & Richard, 1989). Foraging requires animals to use manipulative skills and problem-solving abilities in the process of searching and extracting food items from the environment. Thus, for captive nonhuman primates, providing foraging opportunities is a key component of environmental enrichment that supports species-typical manipulative behavior (Bayne, et al., 1993; Schapiro & Bloomsmith, 1995; Schapiro, 1996). In the captive environment nonhuman primates are given toys or foraging puzzles that require manipulation to extract food and nonfood forage. Evidence for the benefit of this enrichment strategy is found in the results of numerous studies that have demonstrated the provision of such interactive opportunities and varied sensory stimulation reduce the occurrence of abnormal behaviors (Lam, et al., 1991; Novak, et al., 1998).

Foraging opportunities are not without risk of problems, however. In particular, one specific potential negative outcome identified in recent studies is the potential for increased body weight as a function of increased caloric intake via foods used in environmental enrichment. Increases in the body weight of captive animals have been observed over time in a number of different species (Klimentidis, et al., 2011) is a common concern in veterinary practice (Lund 2005; Lund 2006; German, 2006; for review see Bomberg et al. 2017) and weights have increased over time within the nonhuman primate colony at the Wisconsin National Primate Research Center (WNPRC; Terasawa, et al., 2012). However, there is currently no empirical data on the degree to which increased weight of captive nonhuman primates is observed across research facilities. Nor is there direct evidence that increases in body weight are attributable to environmental enrichment. There is, however, a common perception that weight gain and imbalanced nutrition are likely outcomes of environmental enrichment. It is not an unreasonable concern given increasing emphasis on environmental enrichment, expansion of enrichment programs, and the fact that enrichment often involves foods supplemental to the animals’ typical chow-based diet.

In light of increasing evidence about the benefits of foraging opportunities in terms of occupying animals with manipulation and cognitive activities, there is also increasing agreement that best practices in primate care includes provision of more frequent—or more sustained—foraging opportunities. As a result, the frequency of foraging opportunities provided to monkeys in our colony, and likely others, has increased over the past several years to include daily enrichment. It should be noted that foraging opportunities could be increased without adding significant caloric intake beyond the typical diet. For example, in the WNPRC enrichment plan, foraging delivered 1-2 days per week with minimal food presented with “destructible” non-nutritive foraging opportunities (e.g., paper forage). Similar to other facilities fresh fruits or vegetables are delivered as a supplement to chow on non-foraging toy days. Rather than simply placing those in the food hopper, the fruits or vegetables may be prepared or placed in a manner that promotes foraging. At our facility they are placed on the top of the home enclosure in order to increase manipulation and processing time by requiring animals to pull items through the mesh. Increasing foraging opportunities can clearly be accomplished without major increase in food calories. How the appropriate balance is selected is often not considered within the literature and discussions that guide environmental enrichment. In turn, concerns about weight gain, provision of too much sugar, or other types of unbalanced diets, remain unaddressed by any formal method of recording, or systematic consideration within the evolving literature on recommended best practices.

Tension between providing foraging opportunities while minimizing potential negative effects on animals’ health is common topic for dialogue; however, there is a surprising absence of guidelines or common conventions for addressing the concern. Providing animals with preferred foods via environmental enrichment can result in caloric substitution of nutritionally balanced components of the diet. The consumption of nutritionally balanced primate chow, for instance, may be displaced by the availability of more palatable and highly desirable fruits, nuts, vegetables, or even candies. At a superficial level, this concern is addressed by application of the National Research Council guidelines (2003) that suggest a balanced diet from a purified dry chow ranges between 40-110 kcal/kg in adult and 100-200 kcal/kg in juvenile to infant macaques and the recommendation that no more than 5-10% of the diet should be attributable to enrichment foods (Toddes, et al., 1997). To our knowledge, however, few studies or facilities directly assess and report calories provided to animals via chow versus calories delivered from enrichment. Of note, the NRC Guide for the Care of Laboratory Animals (2011)” acknowledges the problems of animals self-selecting more palatable foods (e.g., fruit rather than standard chow), but cite a single study in rodents, not non-human primates (Moore, 1987). Furthermore, there are no widely-used guidelines that could drive decisions about selection of types and portion size of foods used in environmental enrichment. As a result, there is likely variability across facilities that house captive animals. Sources of variability in practices across time and individuals, pose potential challenges to scientific objectives, including threats to replication and to interpretation of results.

Together, consideration of captive animals’ diets is critical not only to scientific objectives, but also to long-term clinical care for the animals. Therefore, more closely monitoring and recording the amount, type, and nutritional profile of food delivered through enrichment practices is important as a refinement that serves scientific objectives and animal care and welfare goals. Many—perhaps even most—facilities do have recommended serving sizes for foods that are supplemental to chow and that are used in environmental enrichment. The recommendations may be based, as they are at WNPRC, on veterinary recommendations (for example, see Table 1). Similarly, the Environmental Enrichment Plan (EEP) of some facilities also specifies the frequency and type of foraging opportunities that are offered to the animals as acknowledgment that some consideration is required in order to provide a diet that is nutritionally balanced.

**Table 1.** WNPRC list of enrichment food items and suggested serving sizes.

| ENRICHMENT ITEM | PORTION SIZE | ENRICHMENT ITEM | PORTION SIZE |
| --- | --- | --- | --- |
| **PRODUCE** |  | **NUTS/SEEDS** |  |
| Apple | 1/4 to 1/3 | Peanuts (unsalted, unshelled) | 1 |
| Banana | 1/4 to 1/3 | Slivered Almonds | NA |
| Cabbage | 1/8 | Sunflower seeds (unshelled) | Small handful |
| Cantaloupe | 1/16 | **OTHER FOOD ITEMS** |  |
| Carrots | 1/3 to 1/2 | Applesauce | 1/2 cup |
| Cauliflower | 2 Florets | Banana Chips | 1 to 2 |
| Clementine | Whole | Cereal – toasted oats, fruit whirls | 1 to 4 |
| Coconut | Whole | Coconut – Shredded | 1/8 - 1/4 cup |
| Cucumbers | 1/4 to 1/3 | Dried Apricots | 1 |
| Eggs, hard-boiled | 1 | Drink mix (sugar-free) | Dilute 50% |
| Grapefruit | 1/6 – 1/8 | Fig Newton | 1/2 to whole |
| Green Beans | 2 to 3 | Fruit Roll-ups | Tear small pieces or whole |
| Green Pepper | 1/4 | Ice Cream Cones | 1 |
| Honeydew Melon | 1/16 | Jelly | ½ Tbsp |
| Kale | 2-3 leaves | Karo Syrup (lite) | 1/2 Tbsp |
| Kiwi | 1/2 | Marshmallows (mini) | 1 to 2 |
| Lemons | 1/4 | Mealworms | NA |
| Lettuce | 1/8 | Oyster Crackers | 3 to 5 |
| Limes | 1/4 | Peanut Butter | 1 Tbsp |
| Oranges | 1/4 | Popcorn ( popped) | 1 cup |
| Papaya | 1/4 | Pumpkin - canned | ½ cup |
| Peaches | 1/2 | Raisins | 1 Tbsp |
| Peapods | 2 to 3 | Tortilla (flour or corn) | 1 |
| Pears | 1/4 | Whole Wheat Bread | 1 slice |
| Pineapple | Small chunk or ½ ring | Yogurt – Strawberry, Vanilla, Plain | 1/2 cup |
| Potato | 1/6 – 1/2 |  |  |
| Pumpkin (seasonal) | 2-4 inch slice |  |  |
| Radishes | 2 to 3 |  |  |
| Spinach | 12-15 leaves |  |  |
| Strawberries | 2 to 3 |  |  |
| Watermelon | Small wedge (~ 1 inch thick) |  |  |
| Zucchini | 1/3 to 1/2 |  |  |

The WNPRC enrichment program includes a rotation of foraging puzzles that have different interactive characteristics and food contents that vary in textural, olfactory, and gustatory qualities. To prepare and deliver our foraging puzzles we employ an integrated team approach. Animal husbandry and behavioral management staff share enrichment duties in order to promote positive human-animal interactions as staff work together to deliver desirable interactive foraging items. Food enrichment is prepared in advance of delivery and the individual preparing the puzzle may not deliver the puzzle to the monkey. The specific puzzles, foods, and serving sizes recommended for each food are part of standard operating procedures for environmental enrichment.

Our goal was to unobtrusively evaluate our current practice in order to identify potential sources of variability in caloric and nutritive content across puzzles, personnel, and food types. We conceptualized our study to not assess compliance or strict adherence to serving size SOPs or activity in a single preparation area, rather we chose to collect data at a timepoint after preparation of foraging puzzle to allow us to evaluate “what the animals actually receive” focusing interest on the variability in puzzle food volume across the entire colony. Further, we aimed to evaluate the extent to which simple records of environmental enrichment delivery (i.e., puzzle type, food type, SOP-specified quantity of food) can serve as a reliable estimate of food delivered to the animals.

The sources of variability in the amount of food that is delivered to the animals do include human factors. For example, a simple scenario involves filling puzzles of different volume to the same relative capacity, a known human tendency (van Kleef, 2012). Thus, caloric load associated with a puzzle would be adjusted up or down based on the puzzle characteristic of volume and could interact with other operational factors, such as, serving size or different food components. Such an outcome would clearly result in the delivery of added calories to the diet with larger volume puzzles. However, the occurrence of overfilled foraging puzzles at a “population” level across the colony is unknown. We chose this colony-wide approach as a first step in the assessment of potential overfilling foraging puzzles.

The broad goals of the study reported here were three-fold: First, to provide quantitative data that can be used to estimate the proportion of nutrition/calories delivered via environmental enrichment; second, to apply the quantitative data to serve direct comparisons that can inform selection of enrichment; third, to develop a straightforward method for these measurements. To do this we: 1) measured the amount and type of food in prepared enrichment puzzles; 2) calculated caloric and nutritional information; 3) assessed variation across puzzle types; and 4) calculated inclusive cost for enrichment puzzles, foods, and labor. To address these goals, we sampled multiple preparation areas, collecting puzzle-loading data after puzzles were prepared. Our facility uses puzzles, foods, and techniques that are common in nonhuman primate enrichment. Therefore, our data likely reflect the range of variation one might expect at other facilities housing nonhuman primates.

## Methods

### Study site

The WNPRC houses two species of macaques (*Macaca mulatta and Macaca fascicularis*). The provisioning of foraging devices in this study focused on our Enrichment plan for macaques. The WNPRC facility cares for approximately 1300 macaques housed at multiple locations. The environmental enrichment puzzles that were the focus of this study have the potential to be delivered to 54 different housing rooms. Animals were either socially-housed in groups of 2 or more monkeys, or singly-housed under a veterinary or research exemption from required social housing. No animals were singly-housed for the purpose of the analysis reported here. The macaques are maintained on a chow-based diet (Harlan-Teklad, diet 2050) and *ad libitum* water. Animals’ body weights are monitored monthly and dietary adjustments are made accordingly to promote healthy growth and minimize fluctuation body weight. Clinical veterinary staff oversees body weight and dietary adjustments. Adult macaques approximate a daily chow intake of a 100 kcal/kg, and adolescents’ intake ranges toward 200 kcal/kg. Animals were treated in accordance with the ethical principles of animal care outlined by US federal regulation and the guidelines described in Guide for the Care and Use of Laboratory Animals (2011).

### Procedure

Animal husbandry and behavioral management staff routinely prepare foraging puzzles for delivery to monkeys by placing food within the puzzle in a manner that allows the monkey to actively manipulate the puzzle to retrieve the food. The puzzles were prepared as part of our EEP and in accordance with WNPRC standard operating procedures (SOPs) for puzzle preparation that prescribe serving size and filling strategy. These SOPs were regularly reviewed and discussed to ensure compliance. The foods were allocated by using recommended serving sizes that are reviewed by the attending veterinarian and are part of the enrichment SOPs (see Table 1).

The general procedure for foraging puzzle delivery requires the staff deliver them to the animals and return 1 to 2 hours later to collect the puzzles for cleaning. All of the puzzles evaluated were hung on the outside of the cages, thus all dropped food would fall out of the animals reach. Many of the prepared puzzles include a liquid or semi-solid medium (yogurt, peanut butter, molasses, etc) to bind fruits, vegetables and dry foods to the puzzle.

Over a 9-month period, puzzle content was measured in five food preparation areas. Approximately 20 individual staff members contributed to puzzle preparation for this study. Puzzles were prepared well in advance of the delivery process (minimally the day before). Just prior to delivery, staff members were trained to record the type of food added and the weight of each prepared puzzle on a digital scale (Ohaus Corporation, Model N1B110). Identical scales were used in each sampling area. The weighing process was straightforward: Staff simply placed the puzzle on the scale and recorded numeric value that the scale displayed. The process was quickly performed, added little burden to enrichment delivery, and produced reliable quantitative data. Ten types of foraging puzzles were evaluated (see Figure 1).

**Figure 1.**
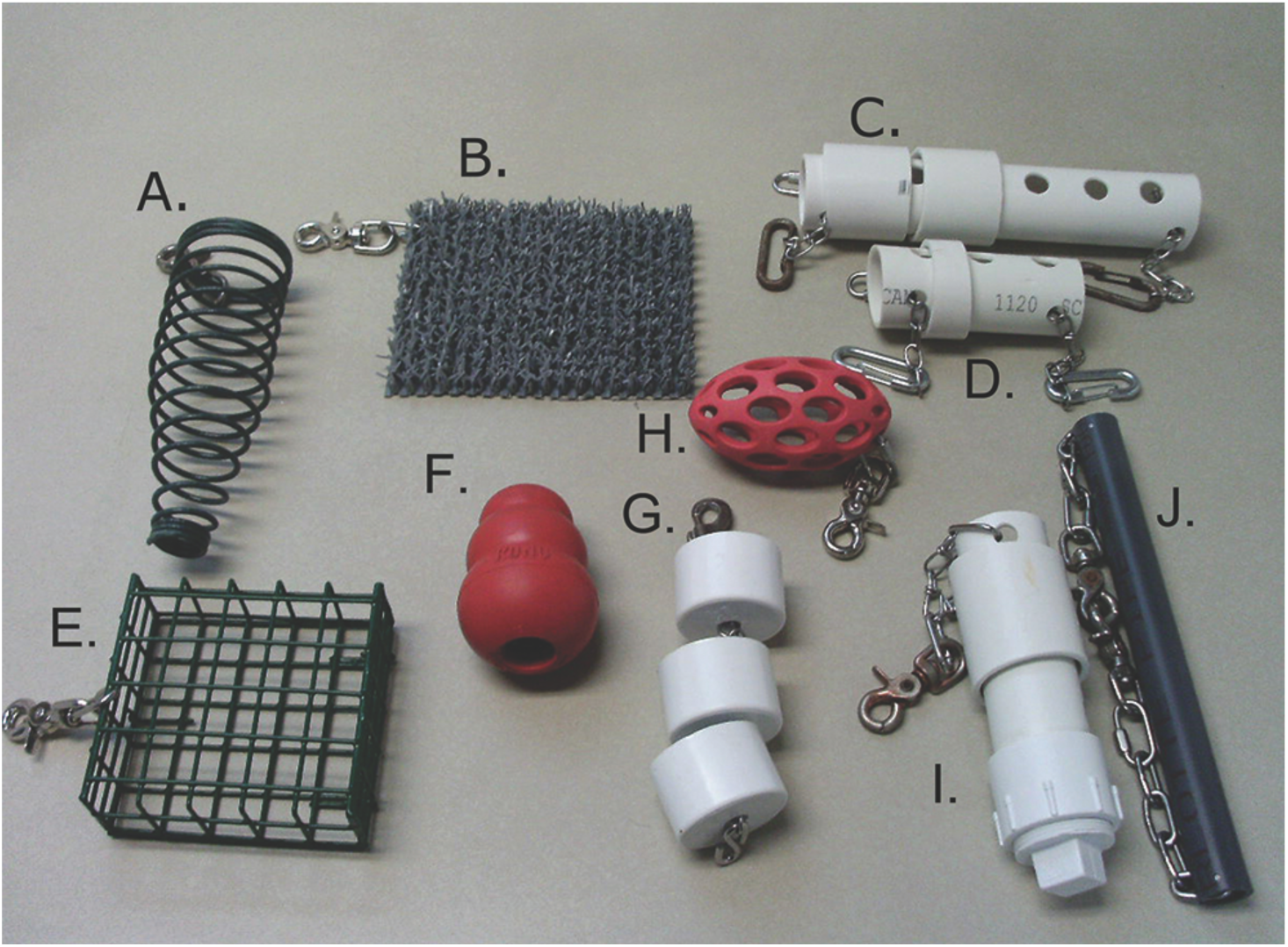
The standard foraging puzzle types sampled for the study: a) Corn feeder, b) foraging turf, c) Large holey sliders, d) small holey slider, e) suet feeder, f) kong, g) cup feeder, h) football, i) peek-a-boo and j) paint roller (top-bottom, left-right).

### Caloric content determination

Our sampling approach recorded the amount of food (total weight) placed in prepared foraging puzzles prior to their delivery to the monkeys. This approach allowed us to record puzzle weights in the absence of the preparer, and thus, to efficiently capture the uncensored amount of food foraging normally delivered to the monkeys.

Measurements occurred as follows: 1) Empty puzzle weights were calculated by weighing clean and empty puzzles that had not yet been prepared for delivery to the animals. For each of the 10 puzzle types, 10 empty puzzles of each type were weighed to determine the average empty puzzle weight. 2) The empty weight was then used to calculate the weight of food in prepared puzzles (i.e., weight of prepared puzzle – weight of empty puzzle = weight of food; see Table 2). At the time of weighing the loaded puzzles, a description of the type and number of items constituting the food component were recorded. For our illustration of caloric load, the gram weight of the standard serving sizes of each food component was calculated (see Table 1 for items and serving size). Each food item contributed as a proportion of the serving size weight to the total load weight. In this manner, the heavier items contribute more calories than the lighter items.

**Table 2:** Characteristics of the ten foraging puzzles sampled for direct comparison.

| Puzzle | Volume (cc)<br>/Area cm<sup>2</sup>* | Observations | Puzzle<br>Characteristic | Min/Max(g) | St Dev. | Median load<br>wt. (g) | Mean load<br>wt. (g) |
| --- | --- | --- | --- | --- | --- | --- | --- |
| Paint Roller<br>(WNPRC) | 231.04* | 25 | Surface | 2.2/87.5 | 24.6 | 51.4 | 46.1 |
| Foraging Turf<br>(Grassworx, LLC/WNPRC) | 252* | 18 | Surface | 12.4/105.8 | 25.8 | 38.5 | 42.9 |
| Football<br>(JW Products, Arlington, TX) | 98.87 | 24 | Internal Vol SM | 10.2/81.8 | 19.4 | 57.3 | 56.0 |
| Small Holey Slider<br>(WNPRC) | 131.1 | 15 | Internal Vol SM | 3.6/112.1 | 33.1 | 58.8 | 54.7 |
| Kong (Medium)<br>(The Kong Co, Golden, Co) | 92.0 | 19 | Internal Vol SM | 43.9/92.0 | 11.8 | 67.4 | 68.9 |
| Peek a boo<br>(WNPRC) | 177.3 | 29 | Internal Vol LG | 38.5/159.1 | 33.4 | 98.4 | 96.6 |
| Cup Feeder<br>(WNPRC) | 168.0 | 30 | Internal Vol LG | 41.2/161.8 | 32.7 | 113.8 | 110.7 |
| Big Holey Slider<br>(WNPRC) | 247.2 | 15 | Internal Vol LG | 12.8/207.9 | 58.1 | 103.4 | 107.4 |
| Suet Feeder<br>(Stokes Seeds, Buffalo, NY) | 576.0 | 24 | Wire | 41.9/178.9 | 38.7 | 139.2 | 122.3 |
| Corn Feeder<br>(Stokes Seeds, Buffalo, NY) | 504.8 | 20 | Wire | 73.2/280.4 | 50.9 | 164.0 | 163.5 |

Note- The characteristics of the ten foraging puzzles sampled. Measures included the volume or area; the number of observations recorded, and puzzle characteristic, minimum and maximum grams contained, standard deviation, median and mean weight in grams for each puzzle type.

The assumption of proportionality of the food components applied to a puzzle serves as a starting point in estimating the calories associated with different foraging puzzles. The strategy of assuming proportionality was chosen over deconstructing food component of individual puzzles for two reasons: 1) our aim was to acquire the data without affecting normal preparation and delivery routines; 2) we aimed to derive estimates broadly from across the entire colony.

To illustrate the potential for serving sizes to increase the calories delivered we extrapolated “standard” and maximum load weights for small and large foraging puzzles based on the weight measurement of prepared puzzles. Standard load was defined as total gram weight of all components of foraging food mass (see Table 2). Thus, standard load weight approximated the mean load weight (g) and maximum load weight approximated the maximum values recorded for small and large foraging puzzles.

Food component serving size weights were used to calculate the caloric values for each food item (Table 3, column 5: Calories (kcal) est.). Serving size weights were used because this is the standard used by the USDA National Nutrient Database for Standard Reference (http://ndb.nal.usda.gov/hdb/foods/list). In this manner, the calculations of load weights and proportional calorie estimates are as follows (Table 3):

**Table 3:**
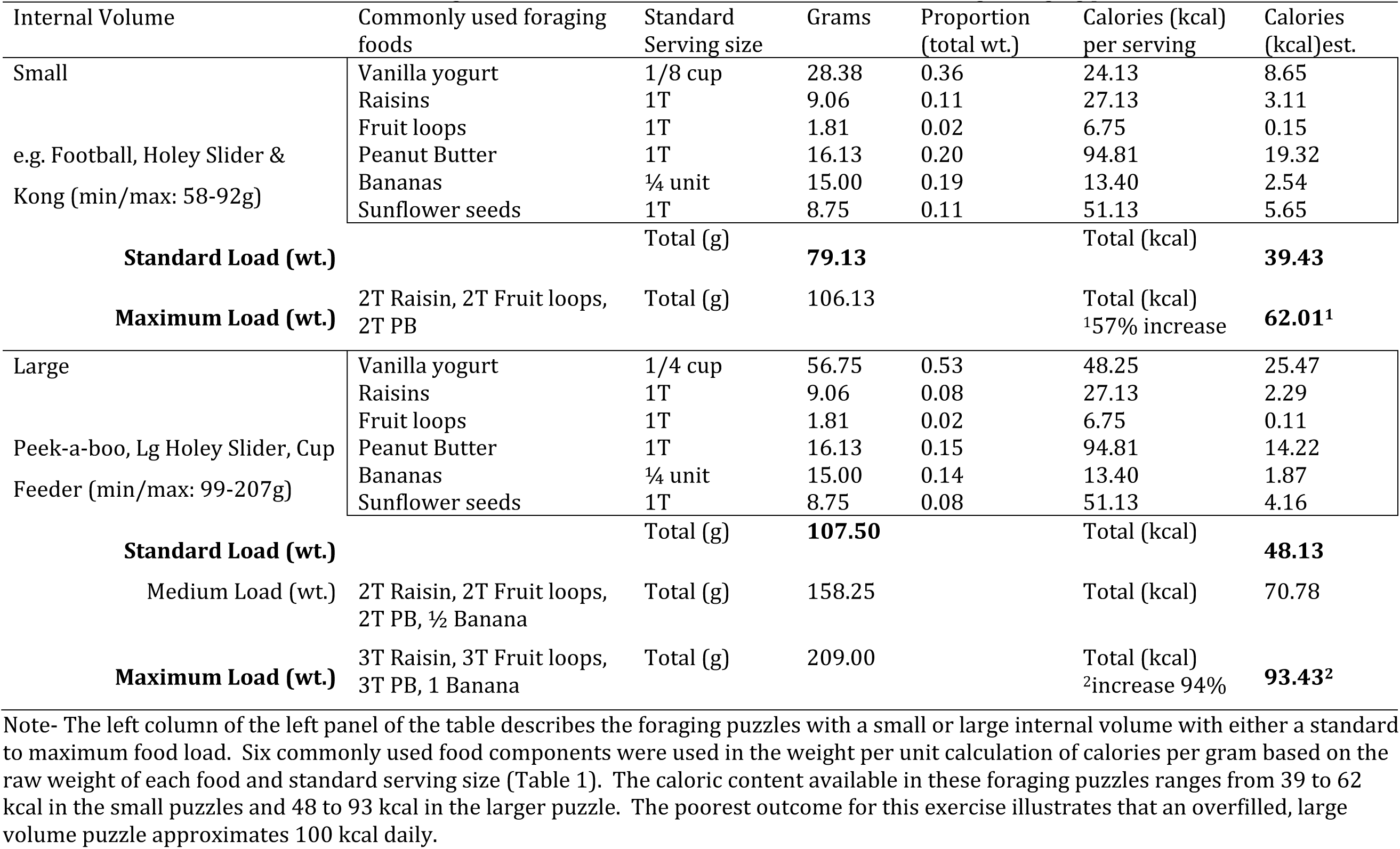
An illustration of the effect of serving size on the caloric content available on small vs. large foraging puzzles.

**Standard load wt.(g)** = Serving size food item 1 (g) + Serving size food item 2 (g) + Serving size food item 3 (g) + Serving size food item *i* (g)*…*

**Proportion of food item to load wt.** = Individual food item (g) /load weight

**Calorie estimation** = kcal per serving * proportion of food item.

### Puzzle Comparisons

Comparisons between puzzles were made in one of two ways: 1) Illustrative comparison of small or large volume puzzles based on puzzle characteristics of volume or surface area (see Table 2); and 2) A direct comparison of two large volume puzzles with similar food components (see Table 4)., foraging turf and cup feeder, we sampled a puzzle of each type from the same preparation area. Our colony employs a mixed-food strategy in which several food items may be used to fill a single puzzle, therefore, it was necessary that the puzzles contain similar foods.

**Table 4.** Illustrative comparison of nutritive components of the food applied to two foraging puzzles.

| Puzzle | Volume (cc)<br>/Area* (cm<sup>2</sup>) | (g)<br>Delivered | Total fat<br>(g) | Total<br>Calories<br>(Kcal) | Total<br>Carbohydrate<br>(g) | Fiber (g) | Sugar (g) | Protein (g) |
| --- | --- | --- | --- | --- | --- | --- | --- | --- |
| Foraging Turf*,<sup>1</sup> | 252.0 | 105.8 | **1.0**<sup>3</sup> | 54.5 | 10.9 | .70 | 8.03 | 1.36 |
| Cup Feeder<sup>2</sup> | 168.0 | 91.7 | **.55** | 46.28 | 9.79 | .32 | 7.48 | 1.23 |

Note- Comparison of total fat, calories, carbohydrate, fiber, sugar and protein for the food applied to a turf (surface puzzle) and cup feeder (large volume puzzle) depicted in the inset of Figure 3. <sup>1</sup>Food components: vanilla yogurt, oatmeal, shredded coconut, and raisins, fruit loops. <sup>2</sup>Food components: vanilla yogurt, yogurt bites, marshmallows, raisins, and captain crunch. <sup>3</sup>85% increase in fat delivered by the foraging turf as compared to the cup feeder although total calories are nearly equivalent.

For our illustrative comparisons of the calories delivered in “small vs. large” puzzles, we simulated a selected combination of food components commonly loaded into puzzles that would yield weight ranges comparable to assessed puzzles (Table 2, Column Mean Load wt. (g)). A standard food load was based on the recommended serving size for a given food type (see Table 1) and because we used a proportional estimation serving size, the heaviest component (e.g. vanilla yogurt) was adjusted from a standard serving of 1/8 cup to 1/4 cup to illustrate how “overfilling” with a single food component might add to the variation of the load weights (Table 3). Similarly, “medium and maximum food load” was used to illustrate the effect of increasing the number of food components or standard serving sizes. Further, the weight (g) for the simulated “standard and maximum load weights” in Table 3, were calculated to fall within the min/max range of total weight of food applied to the 10 puzzles sampled shown in Table 2.

### Cost Analysis

To determine the inclusive cost of our food foraging strategies we used the same method we previously developed and used to assess a range of environmental enrichment strategies (Bennett, et al., 2010; 2014; 2016). In brief, the inclusive cost comprised the initial cost to purchase or construct each puzzle type and the associated labor to prepare (i.e. load the puzzle), deliver, and pre-clean the puzzle after use in preparation for sanitization. As in our previous work, we did not include the cost of water or cleaning chemicals, nor did we account for labor associated more generally with enrichment (e.g., personnel time for ordering supplies, scheduling enrichment, performing compliance-related record-keeping).

### Labor cost calculations

Cost included the labor cycle of preparing a puzzle, retrieving and then pre-cleaning a puzzle for day-to-day use and maintenance. Puzzles were prepared in advance of delivery. To determine puzzle preparation and pre-cleaning times, observers recorded the time they began and finished performing each action. For example, for preparation of puzzles the start time was recorded, as were the number of puzzles prepared, the food elements used and the time the observers finished. Similarly, for pre-cleaning the time was calculated from when the puzzles were taken from the cages to the time the puzzles were ready to be delivered to the sanitization area. The total duration for each action (preparation or pre-clean) was divided by the number of puzzles handled on each sampling occasion. A minimum of 5 preparation/pre-clean observations were used in calculations for each puzzle (Max=11). Puzzles were sampled across many preparation areas. When multiple technicians contributed to a cleaning or preparation session the resulting time values were divided by the number of technicians.

The labor costs were calculated from the average labor charge for an Animal Research Technician from our facility. The time calculations were based on the average duration to complete each task (Preparation and Pre-cleaning). For example, for each occasion a given puzzle was prepared we calculated the number of puzzles prepared per minute (# items/preparation duration) and averaged this value for all observations of a given puzzle. This mean number of puzzles per min was then used to calculate labor cost per puzzle (Puzzles prep per min X Labor rate/60 min.). The same approach was used to calculate labor cost for pre-cleaning each puzzle type. Labor costs for preparation and pre-cleaning were summed for total labor cost per unit. Some puzzles did not require pre-cleaning which reduced the amount of labor required. We also included a commonly used destructible foraging substrate in the analysis in order to represent the full range of cost decisions based on required for maintenance related to provision of foraging opportunities.

## Results

### Effect of puzzle size on amount of food added during preparation of enrichment

Our results demonstrated smaller puzzles had resulted in the delivery of fewer calories an average mean load weight of 59.8 g and large puzzles had a load weight of 104.9 g (T (108) = 8.81, p<. 05; Table 2). Thus, we found that a larger amount of food was applied to the larger puzzles. At the same time, as shown in Table 2, the range in the volume or area and the fill weights of these commonly used foraging puzzles showed overlap and the food weights associated with large puzzles were more variable. These data suggest that large puzzles were not necessarily differentially overloaded, rather the amount of food placed in both small and large puzzles likely exceeded the recommended serving sizes. Similar comparisons can be drawn for surface and wire puzzles.

Table 3 illustrates the caloric impact of the interaction between food types and puzzle type. The calculations indicate that a seemingly small increase in the serving size of high caloric food item may dramatically increase caloric load. For example, a **standard load wt.** for a small puzzle containing vanilla yogurt (28.38g) + raisins (9.06) + Fruit loops (1.81g) + Peanut butter (16.13 g) + Banana (15 g) + Sunflower seeds (8.75g) = 79.13 g. The **proportion of food item to load wt.** (heavy to light) = vanilla yogurt (.36) + raisins (.11) + Fruit loops (.02) + Peanut butter (.20) + Banana (.19) + Sunflower seeds (.11) = 100. The total **Caloric estimation** for or the sum of the caloric estimate for each food component was 39.43 kcal for a small puzzle.

Increasing the serving by a single tablespoon (T) each of raisins, fruit loops and peanut butter, results in a 57% increase in calories, to 62.0 total kcal, the **maximum load wt**. In a larger volume puzzle, an increase from serving size that is proportional to the increased amount for a small puzzle results in even larger disparity between the caloric content of the standard serving size (48.1 calories) and the calories from a maximum loading of the puzzle (93.4 kcal; See Table 3 bold) nearly doubling the calories delivered (increase of 94 %). Thus, Table 3 illustrates relatively small subjective changes (1 to 2 T) of foraging material can substantially increase calories delivered particularly in large devices.

### Comparison of interaction between food type and puzzle characteristics

Table 4 provides direct comparative analysis of food components loaded into (or on: surface) to two sample puzzles that are commonly used in nonhuman primate enrichment: foraging turf and a cup feeder. The difference between the caloric loads for the foraging turf and the cup feeder were similar, 54.5 and 46.2 kcal, respectively. Caloric differences were driven by composition and nutritive content of the food applied to the puzzles. Food items were not matched across the puzzles because food items needed to meet the affordances of the puzzle to best promote foraging. For example, the turf required a sticky component (e.g., yogurt, peanut butter molasses) to adhere lighter items that will stick to the turf surface (i.e., oatmeal, shredded coconut, raisins and fruity loops).

Table 4 illustrates that nutrient amounts varied based on the combination of foods applied, and in turn, these foods differentially contributed to the nutrient profile – i.e. carbohydrate, fiber, sugar, protein. Although the turf and cup feeder delivered comparable caloric amounts, in this case, the nutritive content delivered varied in total fat 1.0 g vs. 0.55g and fiber 0.70 g vs. 0.32g, respectively. The amount of fat delivered by the turf through the “sticky component” (i.e., peanut butter) represents an 85% increase as compared to cup feeder. In turn, this highlights the potential for puzzle characteristics and the food items necessary to promote foraging with them also affect the nutrient components delivered and further illustrated that a proportional component analysis of nutritive elements may be useful in proactive assessments of puzzles.

### Cost analysis

The inclusive cost of purchase and use of the foraging puzzles is summarized in Table 5. The initial purchase or construction costs ranged from a low of USD$0.51 for a “destructible” – a paper plate, bag, napkin or another single-use forage substrate to a high of USD$45.69 for a “holey slider.” Excluding the single-use destructible, the average puzzle cost was USD$ 27.11. The table also provides qualitative assessment of the different puzzles’ longevity, with some-- like the Kong toy-- that are more robust and require less frequent replacement (and thus lower long-term cost). The amount of labor associated with preparing and maintaining the puzzles from delivery to pre-cleaning the puzzles prior to sanitization ranged from $ 0.21 to $1.46, with a mean of $0.72 excluding destructible items. Unsurprisingly, longer labor times and correspondingly higher labor costs were associated with more complex puzzles and delivery of more food components (i.e., a mix of different foods that required preparation).

**Table 5.**
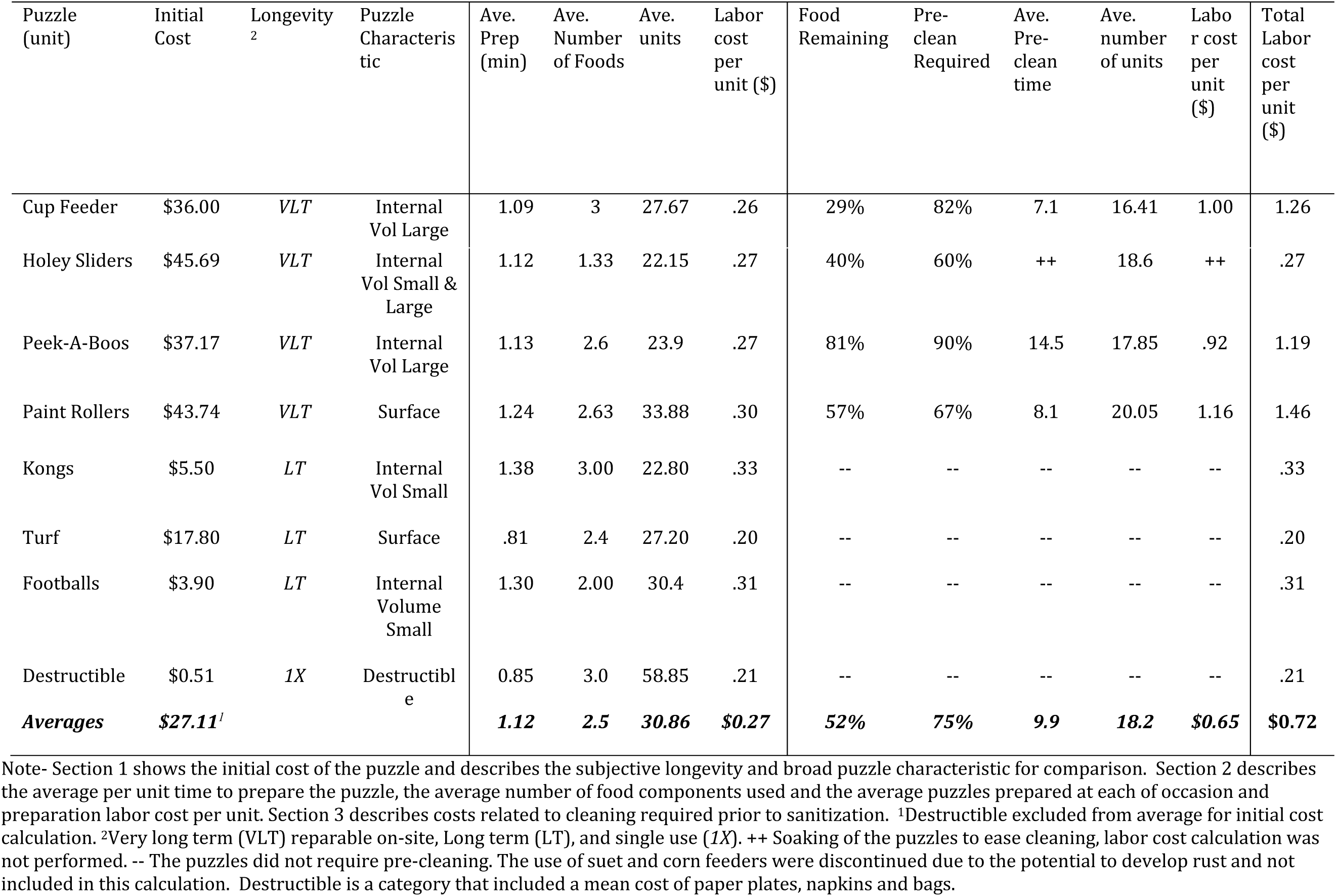
Inclusive cost of purchase and labor associated with foraging puzzle use and maintenance.

| Puzzle (unit) | Initial Cost | Longevity<sup>2</sup> | Puzzle Characteristic | Ave. Prep (min) | Ave. Number of Foods | Ave. units | Labor cost per unit (\$) | Food Remaining | Pre-clean Required | Ave. Pre-clean time | Ave. number of units | Labor cost per unit (\$) | Total Labor cost per unit (\$) |
| --- | --- | --- | --- | --- | --- | --- | --- | --- | --- | --- | --- | --- | --- |
| Cup Feeder | \$36.00 | *VLT* | Internal Vol Large | 1.09 | 3 | 27.67 | .26 | 29% | 82% | 7.1 | 16.41 | 1.00 | 1.26 |
| Holey Sliders | \$45.69 | *VLT* | Internal Vol Small & Large | 1.12 | 1.33 | 22.15 | .27 | 40% | 60% | ++ | 18.6 | ++ | .27 |
| Peek-A-Boos | \$37.17 | *VLT* | Internal Vol Large | 1.13 | 2.6 | 23.9 | .27 | 81% | 90% | 14.5 | 17.85 | .92 | 1.19 |
| Paint Rollers | \$43.74 | *VLT* | Surface | 1.24 | 2.63 | 33.88 | .30 | 57% | 67% | 8.1 | 20.05 | 1.16 | 1.46 |
| Kongs | \$5.50 | *LT* | Internal Vol Small | 1.38 | 3.00 | 22.80 | .33 | -- | -- | -- | -- | -- | .33 |
| Turf | \$17.80 | *LT* | Surface | .81 | 2.4 | 27.20 | .20 | -- | -- | -- | -- | -- | .20 |
| Footballs | \$3.90 | *LT* | Internal Volume Small | 1.30 | 2.00 | 30.4 | .31 | -- | -- | -- | -- | -- | .31 |
| Destructible | \$0.51 | *1X* | Destructible | 0.85 | 3.0 | 58.85 | .21 | -- | -- | -- | -- | -- | .21 |
| **Averages** | **\$27.11<sup>1</sup>** | | | **1.12** | **2.5** | **30.86** | **\$0.27** | **52%** | **75%** | **9.9** | **18.2** | **\$0.65** | **\$0.72** |

Note- Section 1 shows the initial cost of the puzzle and describes the subjective longevity and broad puzzle characteristic for comparison. Section 2 describes the average per unit time to prepare the puzzle, the average number of food components used and the average puzzles prepared at each of occasion and preparation labor cost per unit. Section 3 describes costs related to cleaning required prior to sanitization. <sup>1</sup>Destructible excluded from average for initial cost calculation. <sup>2</sup>Very long term (VLT) reparable on-site, Long term (LT), and single use (1X). ++ Soaking of the puzzles to ease cleaning, labor cost calculation was not performed. -- The puzzles did not require pre-cleaning. The use of suet and corn feeders were discontinued due to the potential to develop rust and not included in this calculation. Destructible is a category that included a mean cost of paper plates, napkins and bags.

## Discussion

The purpose of the foraging component of the enrichment program is to give animals the opportunity to engage in manipulative species-typical behavior. In turn, this is thought to increase the well-being of the animal in the captive environment (Lam, et al., 1991; Novak & Suomi, 1988; Schapiro, et al., 1996; Wolfle, 1999). In general, primate facilities maintain macaques on a chow-based diet to provide for balanced dietary requirements, varying slightly in terms of meal size, pattern, and delivery. Captive-housed macaques receive the bulk of the daily diet in the form of NHP chow, plus any dietary content derived from food enrichment and supplemental fruits and vegetables. The average daily caloric intake derived from chow for an adult macaque in our colony approximates 100 kcal/kg per day (NCR, 2003).

As is common practice in laboratory enrichment, our standard operating procedures include suggested serving sizes to control caloric intake (Table 1). Currently, the caloric or nutritive content of enrichment foods are often not included in diet calculations and the guidelines for the additional contribution related to enrichment or foraging are vague often designating simply that the caloric load should be healthful to the animal. Thus, it is not uncommon for various stakeholders to express concern that “monkeys are getting fat because of enrichment”. Enrichment and enhancements certainly have increased over time, and it is not uncommon to observe individual differences in how puzzles are loaded or to witness an over loaded puzzle. Because most food enrichment appears to be more palatable as compared to chow, it is easy to opine that food enrichment is likely preferred. Further, accommodating accurate serving size is of particular importance given that humans have a bias to overestimate serving size when larger serving bowls are available (see van Kleef, 2012), the unintended consequence of which may negatively affect the animals’ well-being we mean to improve. The reason that this well-known bias is important to evaluate is that it may result in a reapportionment of the “well-balanced” chow-based diet with other enrichment-based foods.

Estimation of calorie and nutrient value is presumed to be one of the central considerations in decisions about foods and puzzles selected to provide foraging opportunities in accord with the NRC recommendation that calories from enrichment should remain within 10-20% of animals’ daily caloric intake (2003). However, there is an absence of quantitative data in the published literature concerning the proportion of caloric and/or nutritive intake delivered through enrichment practices pointing to a potential need for a method to systematic evaluation.

In this study we developed an approach to address this need by using weight and food-type data to provide easily calculated and generalizable estimates of the caloric content of food-puzzle combinations. Our method provides a direct comparison between puzzles based on common characteristics (size) and cost considerations related to use and maintenance. We chose to weigh previously prepared puzzles. To our thinking this removed any potential changes in caretakers’ “puzzle loading behavior” related to the perception that our outcome measures were related to compliance with the standard preparation procedure. The method allowed us to measure puzzles “colony-wide” over all of our food preparation areas and in doing so we were able to assess natural variability in the loaded mass of the foraging puzzles.

It is true that exact calorie counts for each food component could be obtained prior to loading the foraging puzzle. The process for doing so would be to first measure the amount of each food placed in each puzzle and the to calculate the total calories by referring to standard USDA food content data (the same data we use here). Such an endeavor would have required a substantial amount of labor unavailable to us and would not reflect “in situ” loading values. Thus, it should be acknowledged our method is one of many potential approaches and represents an initial step in the development of quantitative measures of dietary calories that can be applied to refining selection and deployment of foraging strategies.

Measuring the weight of prepared food foraging puzzles allowed us to evaluate the relative variability in load size in relation to broad puzzle characteristics (volume or surface area). Table 2 outlines this variation and it is clear that larger puzzles hold more food. However, overlapping variability (Min/max and St dev.) was shown across all of the puzzles sampled which suggests puzzles are not consistently over filled, and thus, reasonably follow the recommended serving sizes for mixed forage. Although the larger puzzles have the potential to hold much more volume it is not the case that large puzzles were always heavier. Further, there was food remaining in many of the puzzles that required extractive action (i.e. cup feeders, holey sliders peek-a-boos and paint rollers) showing that all of the food and calories were not consumed (see Table 5). We also know there is a substantial amount of food waste that falls out of reach with foraging puzzles. For example, the two highest volume puzzles, wire feeders (suet and corn) hang on the outside of the cage front (e.g., see Figure 2). A final point of interest was that the commercially available puzzles do not show less variance in min/max gram range as one might expect if the WNPRC constructed puzzles varied widely in construction tolerances. These observations suggest the amount of food available to collect from these puzzles is both less than the load volume, and likely less than, or within the range of, the suggested serving sizes for food components.

The study illustrates the potential for unintended overfilling of larger volume puzzles. Thus, changes in the number of foraging opportunities with large foraging puzzles may have long-term deleterious consequences. Larger volume puzzles are particularly susceptible to over filling, and therefore, both caloric and nutritive content of food should be closely considered as this may negatively affect both scientific and clinical outcomes (e.g., alter age of pubertal onset and increase diabetes risk; see Cooper, 2015; Klimentidis, et al., 2011, and Terasawa, et al., 2012).

Whether deviation from the actual amount of food delivered and the recommended serving sizes are meaningfully relevant to animal welfare is the question most pertinent to practice and policy decisions. This question depends largely on caloric and nutritional assessment. Table 3 illustrates how increased load sizes can affect the calories made available to the animals through foraging puzzles. To demonstrate we used foods most frequently used in nonhuman primate enrichment programs to simulate a “standard load” based upon mean load weights for the data collected. For example, simply by increasing raisin and peanut butter by a 1T can readily increase caloric content (see bottom of Table 3); however, practical consideration of how puzzles were delivered (i. e., 1 to 2 times per week, rotating puzzles) suggests that it would require the rare case of consistent overfilling of large puzzles over a long period of time to engender weight gain.

Table 2 further illustrates the effect of food loads through comparison of foraging turf and cup feeder (see Figure 2). These two large volume/area puzzles, sampled from the same preparation area with similar foods showed that the estimated calories associated with these puzzles were comparable. The puzzles with a large load size (see Table 3, grams delivered) would contribute an additional 5% caloric content to the diet on any given day. For example, a 10 kg monkey fed a 100 Kcal/kg diet would receive 1000 Kcal per day; the large volume puzzles were estimated to deliver approximately an additional 50 kcal (5 %). This low percentage of additional daily calories falls well within the suggested range of 10-15% (see National Research Council, 2003; Toddes, et al., 1997). However, this percentage would be reduced because all of the food was not attainable (e.g. forage turf), puzzle rotation intersperses large and small volume puzzles, includes food and non-food (e.g., ice and paper forage) contents and the delivery 1-2 per week.

Similar to concerns about caloric input, each of the food items have different nutritive compositions and, while an item may not dramatically increase calories, it may affect values for other nutritive elements. For example, the comparison of fat and fiber applied to the turf and cup feeder in Table 3 provides a rubric by which other investigators can evaluate the food content of the foraging puzzles. Inclusion of other nutritive components is provided in Table 4, and the nutritive information is readily available for this type of calculation (see USDA National Nutrient Database for Standard Reference. http://ndb.nal.usda.gov/hdb/foods/list).

The final goal of this study was to include a cost analysis of each puzzle as a metric for evaluating the benefits and costs inherent to the reviewed puzzles to inform puzzle choices for an enrichment program. The initial consideration of course is the purchase cost of a puzzle, followed by the longevity and cost to employ any given puzzle. Unfortunately, use and maintenance data is not widely available in published literature. Similarly, there are numerous published articles in a number of species that outline the benefits of foraging strategies to the animals; however, few to our knowledge present information on the cost of employing the puzzles.

Table 5 provides the initial cost of the puzzle, a subjective categorization of longevity, puzzle characteristics, average number of food components and metrics related to preparation and pre-cleaning, such as, percentage of puzzles that have food remaining following take down. Per unit estimates of labor cost was presented for each puzzle based upon average hourly labor rate for our facility. Together the information can be used to make data-driven choices both for initial budget start-up costs when developing an enrichment program or when making operational decisions for an existing program. For example, enrichment coordinators could use the cost information to inform choices based on the labor required to maintain a puzzle. Based on labor cost, if delivering 3 opportunities per week one could choose the cup feeder ($1.26), peek-a-boo ($1.23) and turf ($.25) for a total labor cost of $2.74 per week. To be responsive to labor shortfalls or to reduce labor costs strategies one could replace a higher cost puzzle, i.e. the cup feeder with a destructible ($.21) to reduce weekly cost. However, an additional consideration would be to determine the relationship between cost and sustained use. For example, destructible items are cheap but interaction ends with their imminent destruction, whereas more complex foraging puzzles while initially more expensive require continued interaction to retrieve food elements making the interactive value higher for the more expensive puzzle. Taken together, the calculation of values could be used to project monthly or yearly labor costs based on range of mixed puzzles strategies.

This initial study provides a unique approach to assess the broad scale delivery patterns and illustrates the potential effects of puzzle choices and operations associated with employing foraging puzzles. Our intention was to illustrate methods and examples of how to assess calories and nutritive components available to the animal to better refine our ability to provide foraging opportunities that are interesting and healthy. The assessment of load volume as represented both by the physical volume, grams of food, and estimation of the composition of the food components, suggest that it is unlikely that foraging enrichment can serve as a source of added calories sufficient to spur weigh gain as currently deployed under the procedures at our facility.

There certainly was a range of variation in loading habits, but this effect is controlled to some extent by puzzle rotation (small & large load capacitance) characteristics of the puzzle (small, large, non-nutritive, surface feeder) how puzzles are placed in the environment (cage front vs. inside) and waste or non-retrieved food elements. The range of variation shown in prepared puzzle weights suggests that some overfilling does occur. However, our enrichment schedule also incorporates non-nutritive forms of enrichment during a care week ensuring that spikes in caloric load are infrequent and minimized. Furthermore, we schedule food enrichment deliver times distant from chow provisioning likely maximizing chow consumption and ensuring the bulk of the diet is from a nutritively-balanced source. Therefore, conforming to a serving size standard and scheduled frequency of delivery (1 to 2 times per week) is likely sufficient to ensure the animals are not getting a substantial increase in calories from foraging puzzles. Future studies will be required to address both human adherence to standard filling protocols as well as, the outcomes of training that might ensure greater compliance. However, in this assessment we show the estimated calories delivered by foraging puzzles is <10% daily calories. This suggests compliance efforts would be best allocated to emphasize regular rotation of foraging puzzles and conforming to daily schedules, rather than pursuing the burdensome task of evaluating individuals’ preparation styles.

The current study did not directly measure puzzle interaction, waste and amount of daily chow consumed as in previous studies (Bennett et.al., 2010;2014;2016; Gottlieb, et. al., 2011). Rather we favored a broad assessment of “source” calories from puzzles delivered across the colony. Certainly, other sources of excess calories remain for further evaluation, such as supplemental fruits and vegetables, clinical dietary treatments (drug delivery in food), and token rewards by staff. Similarly, dietary allocation studies are necessary to determine preference between chow and foraging enrichment and the animal’s food preferences in the selection of the calories consumed; however, such studies may not be initially amenable to a colony-wide approach (i.e., labor intensive). Rather, smaller scale empirical studies are needed to investigate these sources of caloric variation associated with the interactive and physical characteristics of foraging puzzles.

One could postulate many strategies to limit caloric availability within the care environment; however, deployment decisions may also have less than desired outcomes. For example, one might limit caloric content from foraging by use the same foods regardless of puzzle type; however, this would come at a cost of limiting a desired aspect of foraging, that is, varying textural, olfactory and gustatory qualities. In reality, the characteristics of different puzzles result in differences in which foods can be used to best promote foraging. Puzzle characteristics or affordances of each puzzle require the animal to use different manipulative strategies to retrieve the food. For example, extractive foraging puzzles, such as, a Peek-A-Boo or Holey Slider, have large volumes, but not all of the food material may be accessed (see Table 5; e.g., see percentage food remaining after the 1-2 hrs.). Similarly, corn and suet feeders have a porous structure and paint rollers and turf require animals to pick off food components (Figure1). In each case, food can be displaced and fall out of reach.

Colony-wide evaluations will continue to serve as a necessary step in identifying points of improvement in current practices from which to propose testable hypotheses with focused empirical studies. Refining foraging puzzle use to include consideration of puzzle characteristics, loading patterns, and sources of waste will be required for a complete accounting of the dietary contribution attributable to foraging puzzles. A strength of our colony-wide sampling approach allowed for the estimate the number of calories associated with foraging puzzles prepared in a manner that was unbiased by the influence of observers following the individuals preparing the puzzles. We provided a broad set of methods and examples to identify and evaluate potential sources of variability in calories available through foraging puzzles and suggest other calorie sources that may be of interest for future evaluation.

This study will aid end users by providing a strategy to identify the most cost-effective puzzles-based cost calculations and inform how a strategy contributes to variation in dietary composition delivered through enrichment practices. In practice, one might choose cheap puzzle that is easy to prepare and wonderful for humans to interact with but of minimal interest to the animal. The ultimate goal of providing foraging enrichment should place a premium on increased or sustained interaction and the costs associated with a puzzle should be secondary. Herein lies the balance, in the absence of empirical data to outline both the behavioral (i.e. animal consumer) and operational (i.e. human laborer) costs to inform our decisions on foraging or other enrichment strategies, we must ask: On what are we basing our determination of the relative effectiveness? The current study clearly lies in the area of operational costs estimation; however, puzzle usage analyses will continue to inform how these foraging puzzles invite and sustain species-typical behavior

## Acknowledgements

We gratefully acknowledge assistance from Christine Alioto, and Parker Tenpas in collection of the puzzle weight data. We thank several members of the WNPRC husbandry staff in the collection of puzzle maintenance data: June Bahr, Shawn Busse, Zack Carter, Sam Emmerich, Mia Gambucci, Josue Guadalupe, Leah Fralick, Stephanie Hoker, Amy Laufenberg, Ashley Pratt, Samantha Rettke, Dane Schalk, Dave Thunstrom, Tashi Topgyal, Sarah Woelffler. We thank Deb Jurmu for labor cost information. We thank Bob Becker for figure proofing and formatting. Supported by NIH grant P51OD011106.

## Notes

### Competing Interest Statement

The authors have declared no competing interest.

